# Fusion pore regulation by Epac2/cAMP controls cargo release during insulin exocytosis

**DOI:** 10.1101/403253

**Authors:** Alenka Gucek, Nikhil R Gandasi, Muhmmad Omar-Hmeadi, Marit Bakke, Stein O. Døskeland, Anders Tengholm, Sebastian Barg

## Abstract

Regulated exocytosis establishes a narrow fusion pore as the initial aqueous connection to the extracellular space, through which small transmitter molecules such as ATP can exit. Co-release of larger peptides and hormones like insulin requires further expansion of the pore. There is evidence that pore expansion is regulated and can fail in type-2 diabetes and neurodegenerative disease. Here we report that the cAMP-sensor Epac2 (Rap-GEF4) controls fusion pore behavior by acutely recruiting two pore-restricting proteins, amisyn and dynamin-1, to the exocytosis site in insulin-secreting beta-cells. cAMP elevation leads to pore expansion and peptide release, but not when Epac2 is inactivated pharmacologically or in Epac2^-/-^ mice. Conversely, overexpression of Epac2 impedes pore expansion. Widely used antidiabetic drugs (GLP-1 agonists and sulfonylureas) activate this pathway and thereby paradoxically restrict hormone release. We conclude that Epac2/cAMP controls fusion pore expansion and thus the balance of hormone and transmitter release during insulin granule exocytosis.

## Introduction

Insulin is secreted from pancreatic β-cells and acts on target tissues such as muscle and liver to regulate blood glucose. Secretion of insulin occurs by regulated exocytosis, whereby secretory granules containing the hormone and other bioactive peptides and small molecules fuse with the plasma membrane. The first aqueous contact between granule lumen and the extracellular space is a narrow fusion pore (upper limit 3 nm ^1^) that is thought to consist of both lipids and proteins ^2,3^. At this stage, the pore acts as a molecular sieve that allows release of small transmitter molecules such as nucleotides and catecholamines, but traps larger cargo ^4–7^. Electrophysiological experiments have shown that the fusion pore is short-lived and flickers between closed and open states, suggesting that mechanisms exist that stabilize this channel-like structure and restrict pore expansion ^6,8–10^. The pore can then expand irreversibly (termed full fusion), which leads to mixing of granule and plasma membrane and release of the bulkier hormone content ^4,5,11^. Alternatively, the pore can close indefinitely to allow the granule to be retrieved, apparently intact, into the cell interior (termed kiss&run or cavicapture) ^4,6,12–14^. Estimates in β-cells suggest that 20-50% of all exocytosis in β-cells are transient kiss&run events that do not lead to insulin release ^4,6^. However, kiss&run exocytosis contributes to local signaling within the islet because smaller granule constituents, such as nucleotides, glutamate or GABA, are released even when the fusion pore does not expand. Within the islet, ATP synchronizes β-cells ^15^, and has both inhibitory^16,17^ and stimulatory ^18^ effects on insulin secretion. Within the islet, ATP suppresses glucagon release from α-cells ^19^, and activates macrophages ^20^. Interstitial GABA leads to tonic GABA-A receptor activation and α-cell proliferation ^21,22^, and glutamate stimulates glucagon secretion ^23^.

Regulation of fusion pore behavior is not understood mechanistically, but several cellular signaling events affect both lifetime and flicker behavior. Elevated cytosolic Ca^2+^ and activation of protein kinase C (PKC) accelerate pore expansion and full fusion ^24,25^, while elevated extracellular Ca^2+^ or cAMP, and PI(4,5)P2 favor longer pore lifetimes and kiss&run exocytosis ^6,9,26–28^. Both myosin and the small GTPase dynamin are involved in fusion pore restriction ^29–33^, and assembly of filamentous actin promotes fusion pore expansion ^34^, suggesting a link to endocytosis and the cytoskeleton. In β-cells of type-2 diabetics, upregulation of amysin leads to decreased insulin secretion because fusion pore expansion is impaired ^35^, and the Parkinson’s related protein α-synuclein promotes fusion pore dilation in chromaffin cells and neurons ^36^, thus providing evidence for altered fusion pore behavior in human disease.

Inadequate insulin secretion in type-2 diabetes (T2D) is treated clinically by two main strategies. First, sulfonylureas (e.g., tolbutamide and glibenclamide) close the K_ATP_ channel by binding to its regulatory subunit SUR1, which leads to increased electrical activity and Ca^2+^-influx that triggers insulin secretion ^37^. Sulfonylureas are given orally and are first line treatment for type-2 diabetes in many countries. Second, activation of the receptor for the incretin hormone glucagon-like peptide 1 (GLP-1) raises cytosolic [cAMP] and thereby increases the propensity of insulin granules to undergo exocytosis. Both peptide agonists of the GLP-1 receptor (e.g., exendin-4) and inhibitors of the GLP-1 peptidase DPP-4 are used clinically for this purpose. The effect of cAMP on exocytosis is mediated by a protein-kinase A (PKA) dependent pathway, and by Epac2, a guanine nucleotide exchange factor for the Ras-like small GTPase Rap ^38^ that is recruited to insulin granule docking sites ^39^. Epac2 is also thought to be activated by sulfonylureas ^40^, which may underlie some of their effects on insulin secretion.

Here we have studied fusion pore regulation in pancreatic β-cells, using high resolution live-cell imaging. We report that activation of Epac2, either through GLP1-R/cAMP signaling or via sulfonylurea, restricts expansion of the insulin granule fusion pore by recruiting dynamin and amisyn to the exocytosis site. Activation of this pathway by two classes of antidiabetic drugs therefore hinders full fusion and insulin release, which is expected to reduce their effectiveness as insulin secretagogues.

## Results

### cAMP-dependent fusion pore restriction is regulated by Epac but not PKA

To monitor single granule exocytosis, human pancreatic β-cells were infected with adenovirus encoding the granule marker NPY-Venus and imaged by TIRF microscopy. Exocytosis was evoked by local application of a solution containing 75 mM K^+^, which leads to rapid depolarization and Ca^2+^ influx. Visually, exocytosis of individual granules occurred with two phenotypes. In the first, termed full fusion, fluorescence of a granule that was stably situated at the plasma membrane suddenly vanished during the stimulation (in most cases within <100 ms; Fig 1a-c, left panels). Since the EGFP label is relatively large (3.7 nm vs 3 nm for insulin monomers) this is interpreted as rapid pore widening that allowed general release of granule cargo. It is likely that most of these events involved the collapse of the granule membrane into the plasma membrane. In the second type, the rapid loss of the granule marker was preceded by an increase in its fluorescence that could last for several seconds (flash events, Fig 1a-c, right panels). We and others have previously shown ^41^ that this reflects neutralization of the acidic granule lumen and dequenching of the EGFP-label, before the labeled cargo is released. Since this neutralization occurs as the result of proton flux through the fusion pore, the fluorescence timecourse of these events can be used to quantitatively study fusion pore behavior.

**Figure 1:**
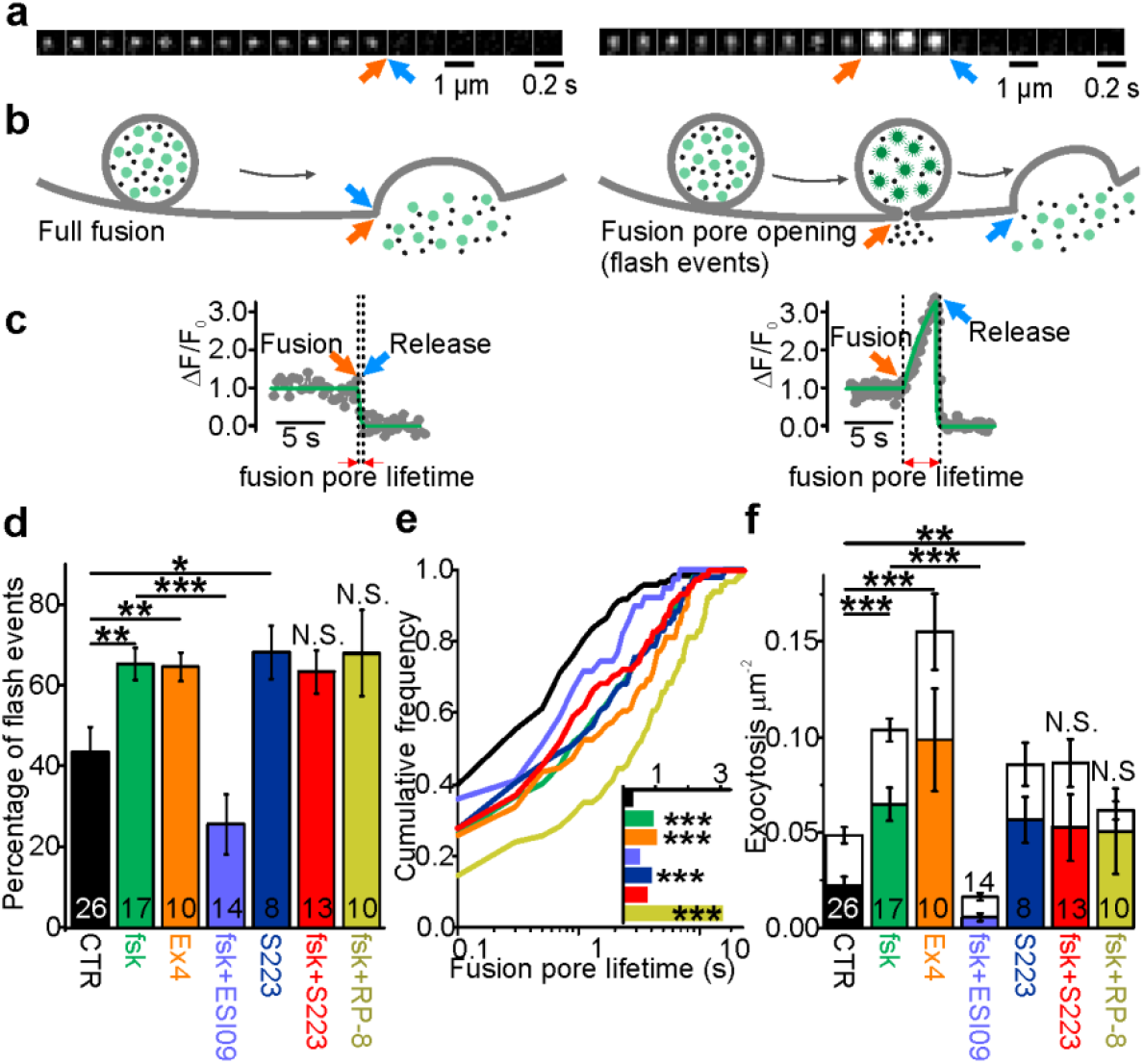
cAMP-dependent fusion pore restriction depends on Epac (but not PKA). **(a)** Examples of single granule exocytosis in human β-cells expressing NPY-Venus and challenged with 75 mM K^+^. Full fusion (left) and flash event (right), where sudden loss of the granule label was preceded by a transient fluorescence increase. Arrows indicate moment of fusion pore opening (orange) and content release (blue). **(b)** Cartoons illustrating the interpretation of events in a. **(c)** Fluorescence time courses for the events in b. Overlaid (green) are fitted functions used to estimate fusion pore lifetime. **(d)** Fraction of flash events in experiments as in a-c, in cells exposed to the indicated agents; forskolin (fsk, P=0.005 vs ctrl), exendin-4 (Ex4, P=0.009 vs ctrl), S223 (P=0.026 vs ctrl), ESI-09 (P=8.5*10^-4^ vs fsk) and fsk+S223 (P=0.74 vs fsk) and RP-8 (P=0.53 vs fsk; u-test). n, number of cells. **(e)** Cumulative frequency histograms of fusion pore lifetimes; fsk (P=8.8*10^-7^ vs ctrl), Ex4 (P=1.9*10^-6^ vs ctrl), S223 (P=6.7*10^-5^ vs ctrl), ESI-09 (P=0.19 vs fsk), Fsk+S223 (P=0.36 vs fsk), and RP-8 P=8.5*10^-5^ vs fsk); Kolmogorov-Smirnov test). Inset shows median lifetimes for 170 (CTR), 197 (fsk), 101 (Ex4), 39 (ESI-09), 94 (S223), 152 (fsk+S223), and 117 (RP-8) events. **(f)** Exocytosis during 40 s of K^+^-stimulation for control (CTR) and with forskolin (fsk, 2 µM, P=5*10^-4^ (P=3*10^-5^ for flashes only), t-test) or Exendin-4 (Ex4, 10 nM, P=1*10^-3^ (P=1.6*10^-4^ for flashes only), t-test) or S223 (5µM, P=0.04) in the bath solution. Inhibitors of Epac (ESI-09, 10 µM, P=2.8*10^-6^ vs fsk (P=1.4*10^-6^ for flashes only), t-test) or PKA (RP-8, 100 µM, n.s. vs fsk, t-test) or Epac2 activator S223 (n.s. vs fsk, t-test) were supplied in addition to forskolin. Flash exocytosis (in color) and full fusions (in white) are shown separately. n, number of cells.

In the following, we will report two parameters that reflect fusion pore behavior, the fraction of exocytosis events with flash phenotype (indicating restricted pores, about 40% in control conditions; Fig 1d), and fusion pore lifetimes estimated by fitting a discontinuous function to the fluorescence timecourse (see Fig 1c, lines and Fig 1e). The distribution of the lifetimes followed a mono-exponential function and was on average 0.87±0.12 s (186 granules in 26 cells) in control conditions (Fig 1e). We limited our analysis to granules that eventually released their peptide content, true kiss&run exocytosis events, corresponding to infinite pore dwell-times. Such events are increased by elevated cAMP ^6,9^ and likely other conditions that stabilize the fusion pore. Indeed, when forskolin (2 µM; fsk) was added to the bath solution we observed a 2-fold increase of exocytosis rate (Fig 1f), a 3-fold increase of fusion pore lifetimes (Fig 1e), and a nearly doubled fraction of events with restricted fusion pores (Fig 1d,f). The GLP-1 agonist exendin-4 (10 nM; Ex4) had comparable effects (Fig 1d-f). Effects similar to those observed for human β-cells (Fig 1) were observed in the insulin secreting cell line INS-1 (Suppl Fig 1).

The effect of fsk on fusion pore behavior was mimicked by the specific Epac2 agonist S223 ^42^. Incubation with S223-acetomethoxyester (5µM) increased the fraction of flash events by 60% (Fig 1d), tripled average lifetimes (Fig 1e) and doubled the event frequency (Fig 1f); the effects of fsk and S223 were not additive. In contrast, the Epac-inhibitor ESI-09 decreased the exocytosis rate in the presence of fsk by 90% (Fig 1f), and the average fusion pore lifetime and the fraction of flash events were both reduced by 60% (Fig 1d-e). PKA inhibition with Rp8-Br-cAMPS ^43^ decreased exocytosis by 40% but not the fraction of flash events, although average fusion pore lifetimes were increased compared with fsk alone (Fig 1e). The results indicate that Epac rather than PKA is responsible for cAMP-dependent fusion pore regulation. Paradoxically, Epac activation increases the rate of exocytosis but slows the rate of peptide release from individual granules.

### Epac2 overexpression restricts fusion pores and prolongs their lifetime

We studied next the effect of Epac2 overexpression on fusion pore regulation. INS-1 cells were co-transfected with EGFP-Epac2 and NPY-tdmOrange2 and fluorescence was recorded simultaneously in both color channels. Epac2 overexpression had no effect on the overall exocytosis rate in either absence or presence of fsk (Fig 2a), but increased the rate of flash events (Fig 2b-c), supporting our finding, based on manipulation of the endogenous Epac2 activity, that Epac2 is involved in fusion pore regulation (Fig 1). Fusion pore lifetimes in cells overexpressing Epac2 increased 3-fold in the absence of fsk, and were similar to controls in presence of fsk (Fig 2d). This indicates that a high Epac concentration can achieve sufficient activity to affect insulin secretion even at basal cAMP level, likely because cAMP acts in part by increasing the Epac concentration at the plasma membrane ^39^.

**Figure 2:**
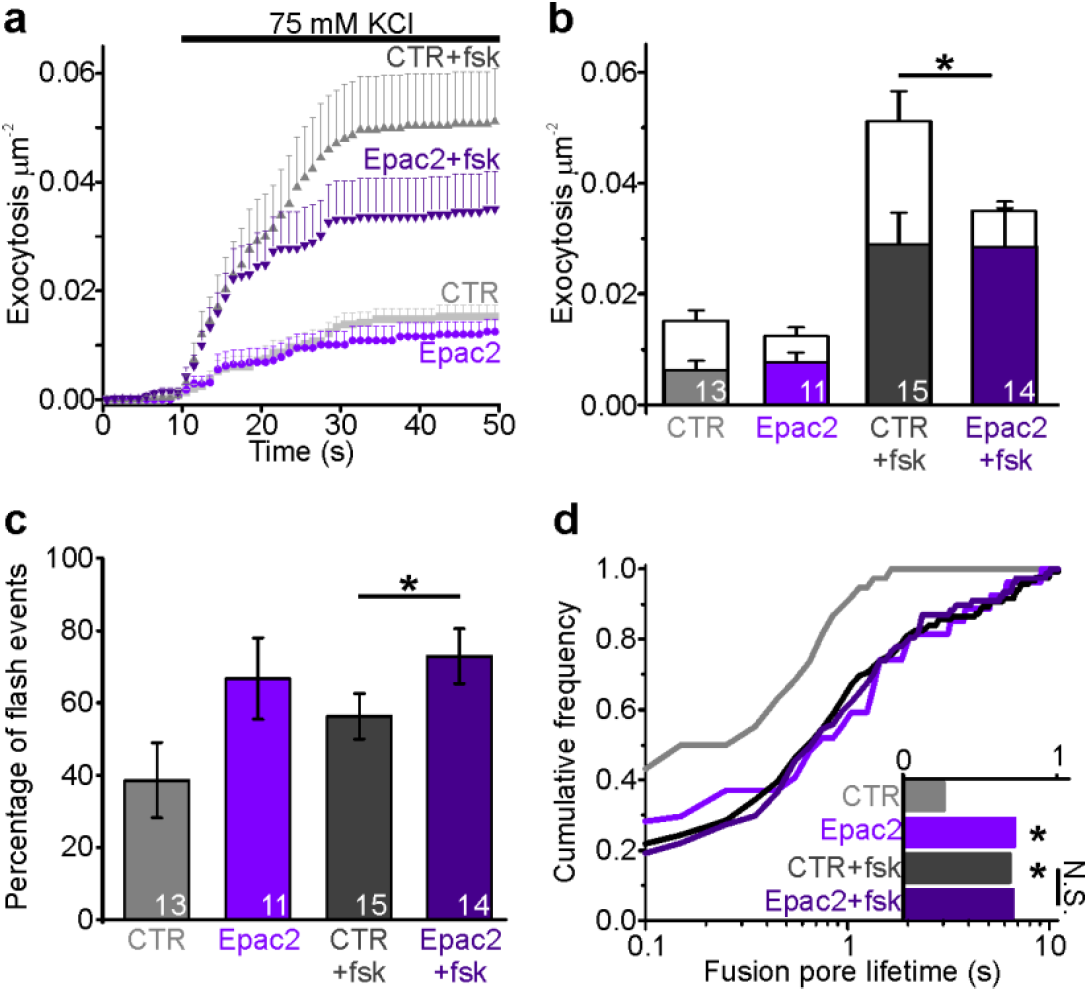
Epac2 overexpression prolongs fusion pore lifetimes. **(a)** Cumulative exocytosis in INS-1 cells stimulated with 75 mM K^+^; grey for control cells, purple for cells expressing Epac2-EGFP (both also expressed NPY-tdmOrange2); fsk indicates forskolin in the bath solution. CTR, n=13; Epac2, n=13; CTR+fsk, n=15; Epac2+fsk, n=14 cells. **(b)** Total exocytosis in (a), separated into full fusion (white) and flash events (color). In presence of fsk, Epac2 expression specifically reduced full fusion events (P=0.011, t-test). n, number of cells. **(c)** Fraction of flash events in (a-b). With fsk, difference was significant (P=0.04, u-test). n, number of cells. **(d)** Fusion pore lifetimes for conditions in a-c. Epac overexpression increased pore lifetimes in absence (P=0.014) but not in presence of fsk (P=0.87, Kolmogorov-Smirnov test). Inset shows the lifetimes for 38 (CTR), 27 (Epac2), 119 (CTR+fsk) and 77 (Epac2+fsk) events.

### ATP release is accelerated upon Epac inhibition

To test if cAMP-dependent fusion pore restriction affects release of small transmitter molecules, we quantified nucleotide release kinetics from individual granules using patch clamp electrophysiology. The purinergic receptor cation channel P2X_2_, tagged with RFP (P2X_2_-RFP), was expressed in INS-1 cells as an autaptic nucleotide sensor (Fig 3a). The cells were voltage-clamped in whole-cell mode and exocytosis was elicited by including a solution with elevated free Ca^2+^ (calculated 600 nM) in the patch electrode. In this configuration, every exocytosis event that co-releases nucleotides causes an inward current spike, similar to those observed by carbon fiber amperometry (Fig 3a-b). Including cAMP in the pipette solution doubled the frequency of current spikes, consistent with accelerated exocytosis. This effect of cAMP was blocked if the Epac inhibitor ESI-09 was present (Fig 3b-c). The current spikes (see Fig 3a, right) reflect nucleotide release kinetics during individual exocytosis events. In the presence of cAMP, but not cAMP+ESI-09, they were markedly widened as indicated by on average 20% longer half-widths (Fig 3d), 30% longer decay constants (τ, Fig 3e), and 40% slower rising phases (25-75% slope, Fig 3f), compared with control. This indicates that nucleotide release is slowed by cAMP, likely because of changed fusion pore kinetics. Since the effect is blocked by ESI-09, we conclude that the cAMP effect probably is mediated by Epac.

**Figure 3:**
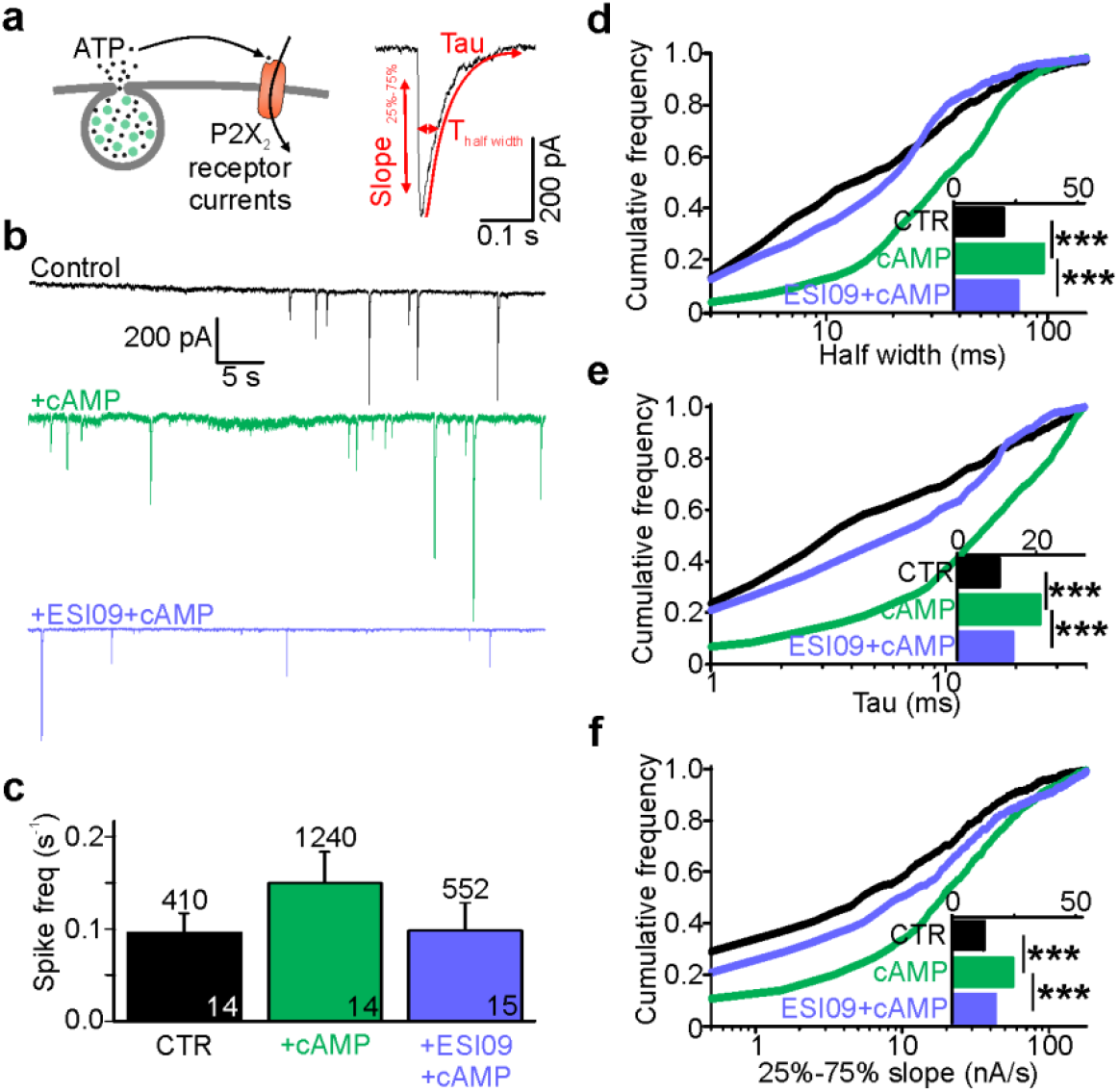
Cytosolic cAMP slows ATP release by activating Epac. **(a)** Electrophysiological detection of nucleotide release events in INS-1 cells expressing P2X_2_-RFP. Cartoon of the assay (left) and example current spike (black) with fit and analysis parameters (red; T_half_, tau and slope during 25% to 75% of peak). **(b)** Representative P2X_2_ currents for control (black), and with cAMP (green) or with cAMP together with ESI-09 (purple) in the electrode solution. **(c)** Spike frequency conditions in (b). n of events (on top) and n of cells (on bars). **(d-f)** Cumulative frequency histograms of spike half width (d), decay constant tau (e), and slope of the rising phase (25% and 75% of peak, (f)) for CTR (n=410 spikes, 14 cells), +cAMP (n=1240, 14 cells) and +ESI-09+cAMP (n=552, 15 cells) with medians in the insets. cAMP increased half-width (P=4.1*10^-31^ vs ctrl, Kolmogorov-Smirnov test), tau (P=2.7*10^-32^, Kolmogorov-Smirnov test), and rising slope (P=4.7*10^-19^, Kolmogorov-Smirnov test); the effects were reversed by ESI-09 (P=3.4*10^-21^, P=3.6*10^-22^, and P=1.3*10^-9^, Kolmogorov-Smirnov test), respectively.

### cAMP-dependent fusion pore regulation is absent in Epac2^-/-^ β-cells

Since ESI-09 blocks all Epac isoforms^44^, we characterized fusion pore behavior in isolated β-cells from Epac2^-/-^ mice, in which all subforms of Epac2 are deleted ^45^. Cells from WT or Epac2^-/-^ mice were infected with adenovirus encoding the granule marker NPY-tdmOrange2 and challenged with 75 mM K^+^ (Fig 4a-b). In the absence of forskolin, exocytosis was significantly slower in Epac2^-/-^ cells than WT cells, and the fraction of flash-associated exocytosis events was five-fold lower (Fig 4c-e). This was paralleled by strikingly shorter fusion pore life-times in Epac2^-/-^ cells compared with WT (Fig 4f). As expected, forskolin increased both exocytosis (Fig 4e) and the fraction of flash events (Fig 4c) of WT cells. In contrast, exocytosis was not accelerated by forskolin in Epac2^-/-^ cells, and the fusion pore lifetimes and fraction of flash events were similar with or without forskolin (Fig 4c, f-g). We conclude therefore that the effects of cAMP on fusion pore behavior are mediated specifically by Epac2.

**Figure 4:**
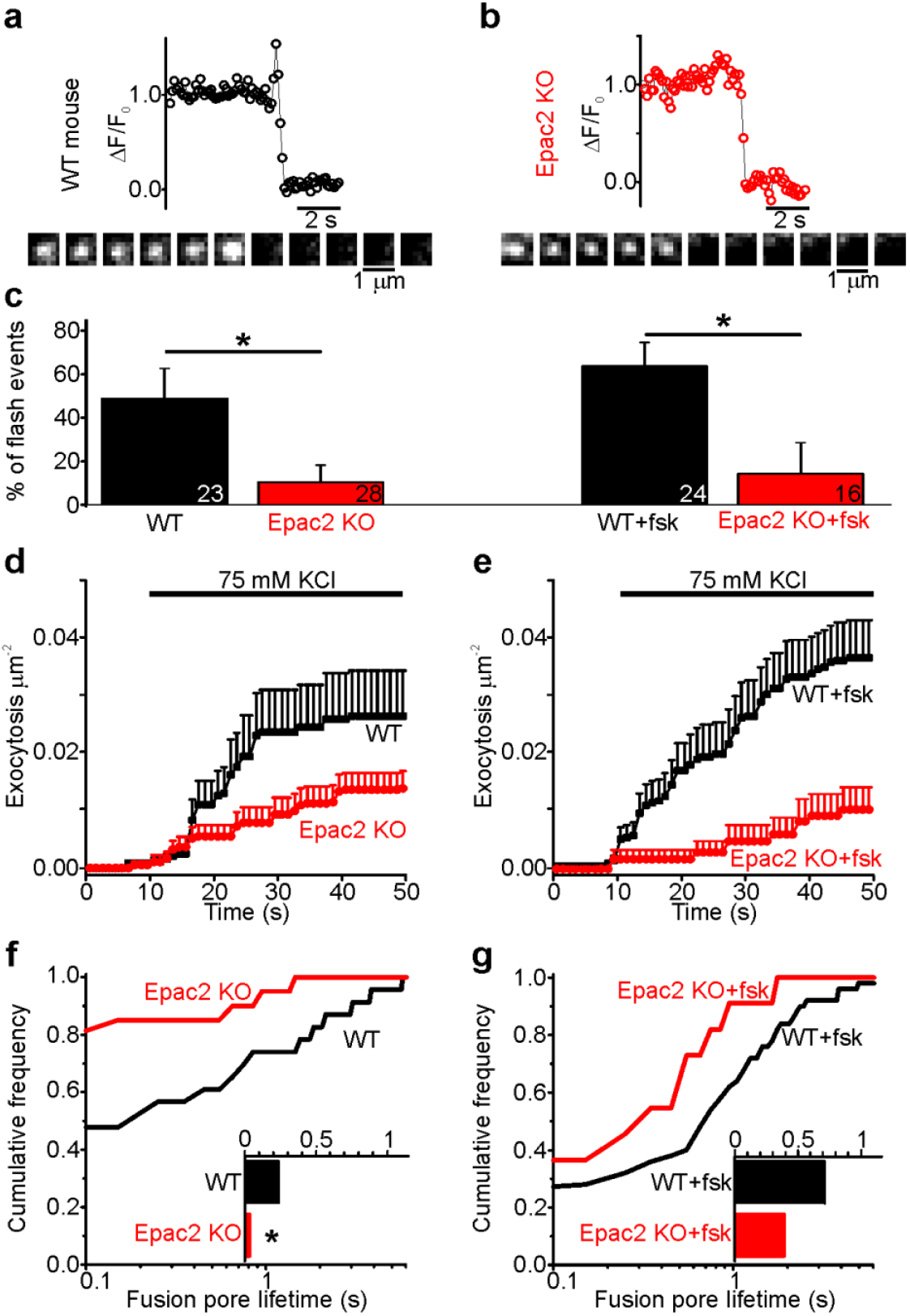
Fusion pores expand rapidly in Epac2^-/-^ mice. **(a-b)** Examples of NPY-tdmOrange2 exocytosis events in β-cells from Epac2^-/-^ mice (b) or from wildtype littermates (a), stimulated with 75 mM K^+^ in presence of forskolin. Note absence of a flash in (b). **(c)** Fraction of flash events for experiments in (a-b). The differences were significant in absence (P=0.028, u-test) or presence of fsk (P=0.017, u-test). n, number of cells. **(d)** Cumulative exocytosis for experiments in absence of forskolin (a,c left) for wildtype (black) and Epac2^-/-^ cells (red).**(e)** Cumulative exocytosis for experiments in presence of forskolin (b,c right) for wildtype (black) and Epac2^-/-^ cells (red). P=0.003, u-test) compared to wt (black). **(f-g)** Cumulative frequency histograms and medians (inset) of fusion pore lifetimes for exocytotic events in d (no forskolin, 23 events for wt, 22 for Epac2^-/-^) and E (with forskolin, 50 events for wt, 9 for Epac2-/-). Differences in f are significant (P=0.043; Kolmogorov-Smirnov test).

### Sulfonylureas delay fusion pore expansion through the same pathway as cAMP

Sulfonylureas have been reported to activate Epac ^40^, in addition to their classical role that involves the sulfonylurea receptor (SUR). We tested therefore if sulfonylureas (SUs) could affect fusion pore behavior. INS-1 cells expressing NPY-EGFP were tested with three types of SUs, with different relative membrane permeability (tolbutamide<glibenclamide<gliclazide). In addition, diazoxide (200 µM) was present to prevent electrical activity. Exocytosis was not observed under these conditions, but could be triggered by local application of elevated K^+^ (75 mM). In the absence of fsk, all three sulfonylureas accelerated K^+^-stimulated exocytosis 2-3-fold over that observed in control (Fig 5b, left), which is consistent with earlier findings that sulfonylureas augment insulin secretion via intracellular targets ^46,47^. This effect was entirely due to an increase in flash-associated exocytosis events (Fig 5b-c) and the average fusion pore lifetime increased accordingly in the presence of sulfonylurea (Fig 5d). In the presence of fsk, which strongly stimulated both flash-associated and full fusion exocytosis in absence of sulfonylurea (Fig 5b-c, middle), the addition of sulfonylureas decreased strongly full-fusion exocytosis, but had no additional effect on the frequency of flash-associated events (Fig 5b, middle). Accordingly, fusion pore lifetimes were elevated compared with control (no fsk), and only marginally longer than with fsk alone (Fig 5d, right). Similar results were obtained in human β-cells, where glibenclamide increased exocytosis in the absence of fsk (P=0.01, n=13 cells) but not in its presence (P=0.80, n=7 cells; data not shown). The data indicate that sulfonylureas restrict fusion pore expansion through the same intracellular pathway as cAMP, which may counteract their stimulating effect on exocytosis by preventing or delaying peptide release.

**Figure 5:**
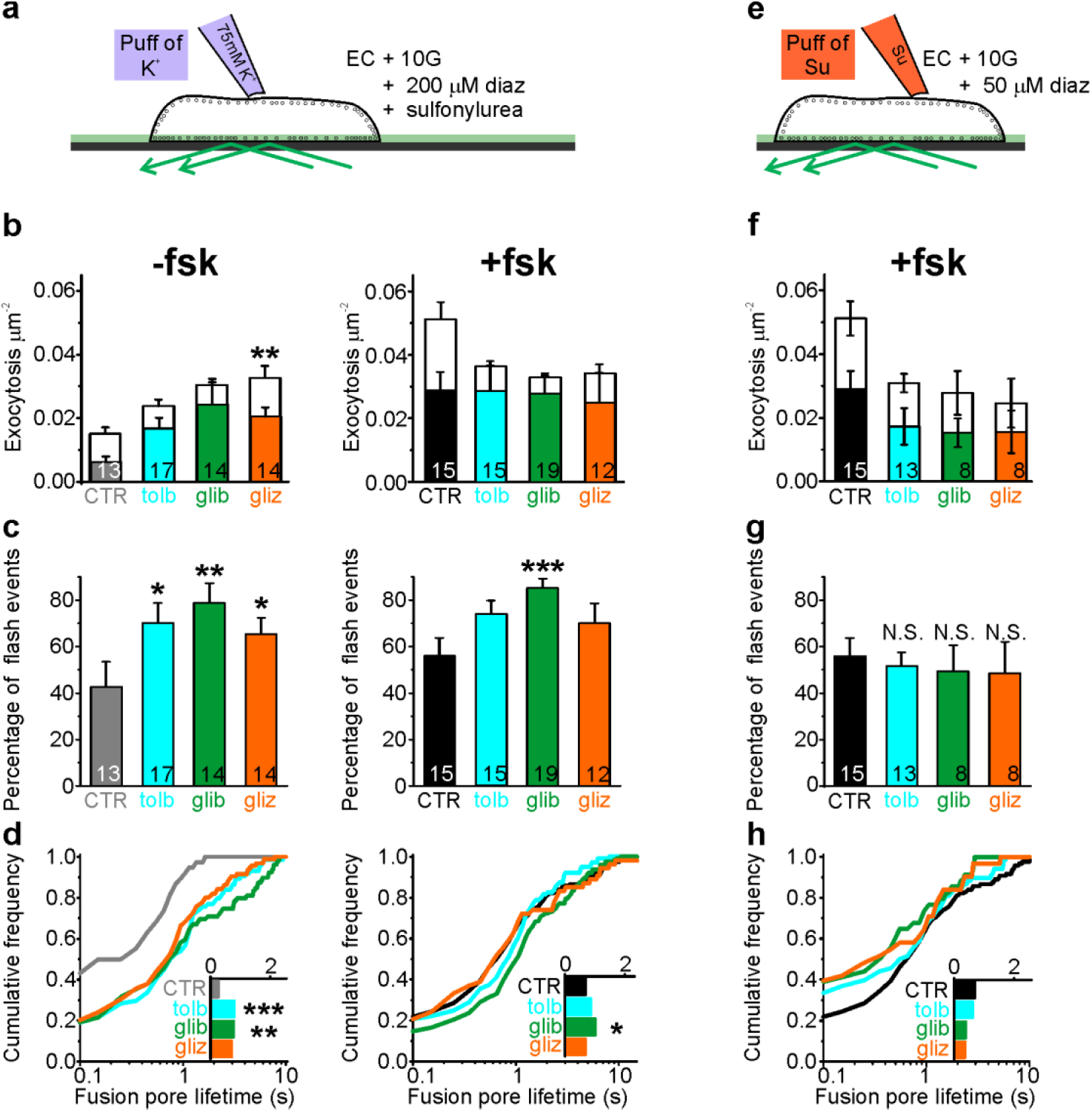
Sulfonylureas cause fusion pore restriction. **(a)** Cartoon of the experimental design in (b-d). INS-1 cells expressing NPY-tdmOrange2 were bathed in 10 mM glucose, diazoxide (200 µM) and either 200 µM tolbutamide (tolb), 50 µM glibenclamide (glib) or 50 µM gliclizide (gliz); exocytosis was evoked by acute exposure to 75 mM K^+^. **(b)** Exocytosis in absence (left) or presence (right) of fsk (2 µM) for flash events (color) and full fusions (white). Total exocytosis was increased by sulfonylurea in absence of fsk (P=0.09 tolb; P=0.07 glib, P=0.003 gliz, t-test), but not in its presence. Sulfonylurea reduced full fusion events in presence of fsk (P=0.0045 tolb, P=0.00032 glib, 0.022 gliz, t-test). n, number of cells. **(c)** Fraction of flash events for experiments in (b); differences vs control are significant (-fsk: P=0.02 tolb, P=0.007 glib, P=0.037 gliz, and +fsk: P=0.05 tolb; P=6.9*10^-4^ glib; P=0.097 gliz, u-test). n, number of cells. **(d)** Cumulative frequency histograms and medians (insets) of fusion pore lifetimes for **(b-c)**. Differences vs control are significant in the absence of fsk: P=9.1*10^-4^ tolb, P=0.003 glib, P=0.015 gliz, Kolmogorov-Smirnov test). Insets show lifetimes for 38 (CTR), 74 (tolb), 79 (glib), 95 (gliz) events and inset on the right for 111 (CTR), 104 (tolb), 127 (glib) and 54 (gliz) events in presence of fsk. **(e)** Cartoon of the experimental design in **(f-h)**. Cells were bathed in 10 mM glucose, 2 µM fsk, 50 µM diazoxide and acutely exposed to sulfonylureas (500 µM tolb, 100 µM glib or 100 µM gliz) during the recording period.**(f)** Exocytosis in presence of fsk (2 µM) for flash events (color) and full fusions (white). Differences in total exocytosis were not significant (P=0.12 tolb, P=0.12 glib, P=0.84 gliz, t-test). n, number of cells. **(g)** Fraction of flash events for experiments in (f). **(h)** Cumulative frequency histograms and medians (inset) of fusion pore lifetimes for (f-g). Inset shows lifetimes for 111 (CTR), 68 (tolb), 34 (glib) and 31 (gliz) events.

To test how sulfonylureas affect exocytosis through direct stimulation of SUR1 and K_ATP_ channels, we applied sulfonylureas also acutely, to avoid their accumulation in the cytosol (Fig 5e). Reduced diazoxide (50 µM) prevented glucose-dependent exocytosis but still allowed acute stimulation of exocytosis by sulfonylureas. Under these conditions, for all three sulfonylureas the fraction of flash-associated exocytosis events (Fig 5f-g) and the fusion pore lifetimes (Fig 5h) was similar to control (stimulation with elevated K^+^). Taken together, the data suggest that sulfonylureas must enter the cytosol to affect fusion pore behavior, and that this effect is not mediated by the plasma membrane SUR. We excluded the possibility that sulfonylureas affect the fluorescence signal indirectly, by altering granule pH (Suppl Fig 2). Moreover, an EGFP-tagged SUR1 (EGFP-SUR1) expressed in INS-1 cells did not localize to exocytosis sites or affect fusion pore behavior (Suppl Fig 3). We therefore conclude that sulfonylureas affect fusion pore behavior through Epac2.

### Dynamin and amisyn-controlled restriction of the fusion pore is cAMP-dependent

The proteins dynamin and amisyn ^30,35^ have previously been implicated in fusion pore regulation in β-cells. To understand how these proteins behave around the release site EGFP-tagged dynamin1 (Fig 6a) or mCherry-tagged amisyn (Fig 6b) were expressed together with a granule marker in INS-1 cells, and exocytosis stimulated with elevated K^+^. In the presence of fsk both of the two fluorescent proteins were recruited to the granule site during membrane fusion (Fig 6c,f), and their overexpression markedly prolonged the fusion pore lifetimes (Fig 6d;g) and increased the fraction of flash-associated exocytosis events (Fig 6e,h). In the absence of fsk, overexpression of the two proteins had no effect on fusion pore behavior and amisyn (but not dynamin1) was recruited to the exocytosis site (Fig 6i-n). The data suggest that dynamin1 and amisyn are acutely recruited to the exocytosis site, where they contribute to fusion pore restriction in a cAMP-dependent manner.

**Figure 6:**
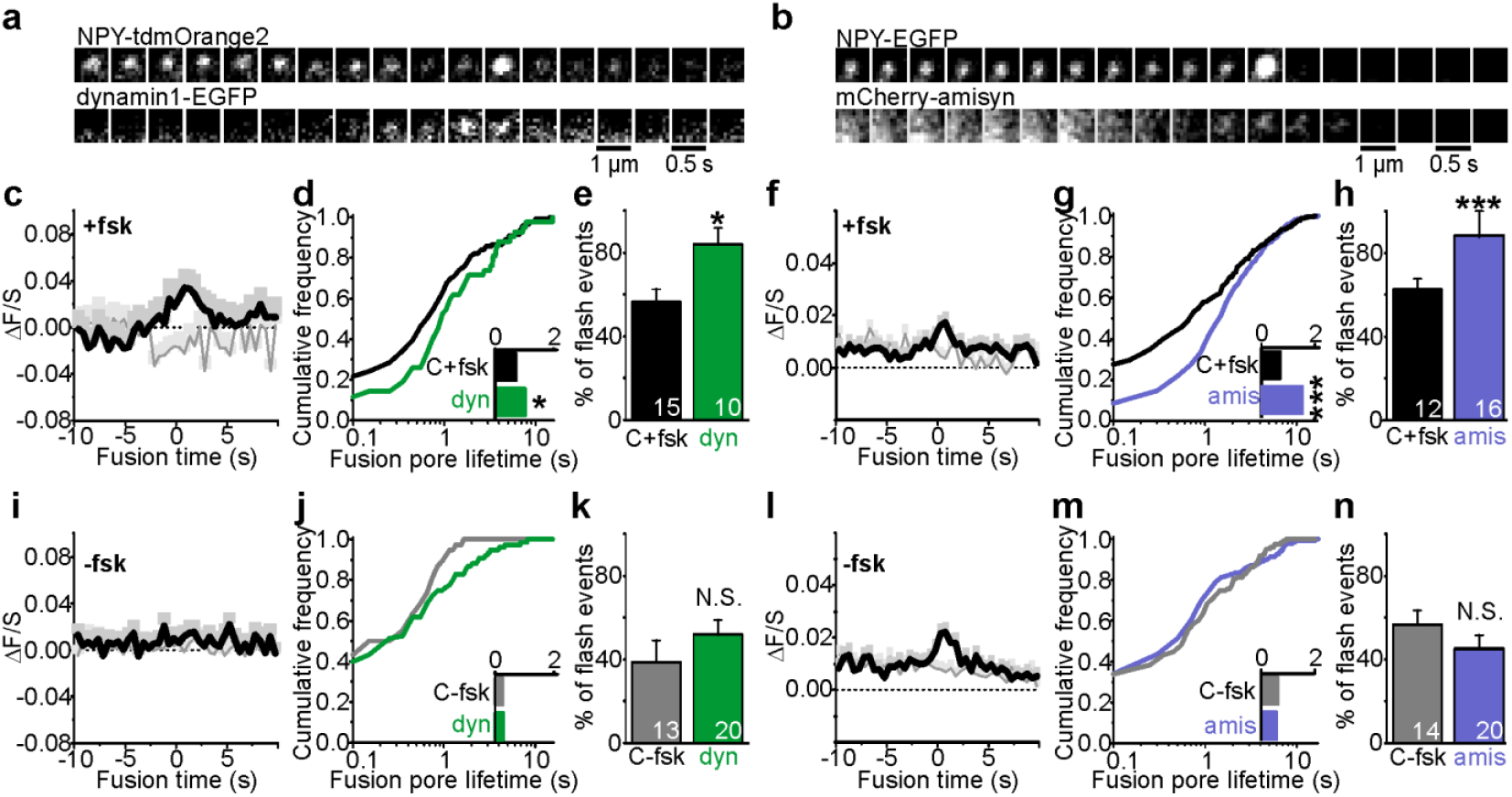
Fusion pore regulation by dynamin and amisyn is cAMP-dependent. **(a-b)** Examples of transient recruitment of dynamin1-EGFP (a, lower) or mCherry-amisyn (b, lower) to granules (upper, labeled with NPY-tdmOrange2 or NPY-EGFP) during K^+^-stimulated exocytosis in presence of forskolin.**(c)** Average time course (±SEM) of dynamin1-EGFP fluorescence during 34 flash-type exocytosis events (in black) and 8 full-fusion type events (in gray) in presence of forskolin; time relative to the onset of the flash in the granule signal.**(d)** Cumulative frequency histograms and medians (inset) of fusion pore lifetimes in cells expressing dynamin1-EGFP (green) or control; fsk was present (P=0.03, Kolmogorov-Smirnov test). 119 (CTR) and 42 (dynamin1) events.**(e)** Fraction of flash events in (d); P=0.016, u-test. n, number of cells.**(f)** Average time course (±SEM) of mCherry-amisyn fluorescence (n=274 flash events, n=46 full fusion events).**(g)** Cumulative frequency histograms and medians (inset) of fusion pore lifetimes in cells expressing mCherry-amisyn (purple) or control; fsk was present (P=2.3*10^-80^, Kolmogorov-Smirnov test). 213 (CTR) and 320 (amisyn) events. **(h)** Fraction of flash events in (g); P=6.3E-4, u-test. n, number of cells. **(i)** As in c, but without forskolin present; n=37 flash events, n=39 full fusion events. **(j-k)** As in (d-e), but for 38 (CTR) and 76 (dynamin1) events in the absence of forskolin **(l)** As in f, but without forskolin present; n=65 flash events, n=73 full fusion events. **(m-n)** As in (g-h), but for 123 (CTR) and 138 (amisyn) events in the absence of forskolin

## Discussion

cAMP-dependent signaling restricts fusion pore expansion and promotes kiss&run exocytosis in β-cells ^9^ and neuroendocrine cells ^27,48^ (but see ^49^). We show here that the cAMP-mediator Epac2 orchestrates these effects by engaging dynamin and perhaps other endocytosis-related proteins at the release site (Fig 7). Since the fusion pore acts as a molecular sieve, the consequence is that insulin and other peptides remain trapped within the granule, while smaller transmitter molecules with para- or autocrine function are released ^4,6,7,13,50–52^. Incretin signaling and Epac activation therefore delays, or altogether prevents insulin secretion from individual granules, while promoting paracrine intra-islet communication that is based mostly on release of small transmitter molecules.

**Figure 7:**
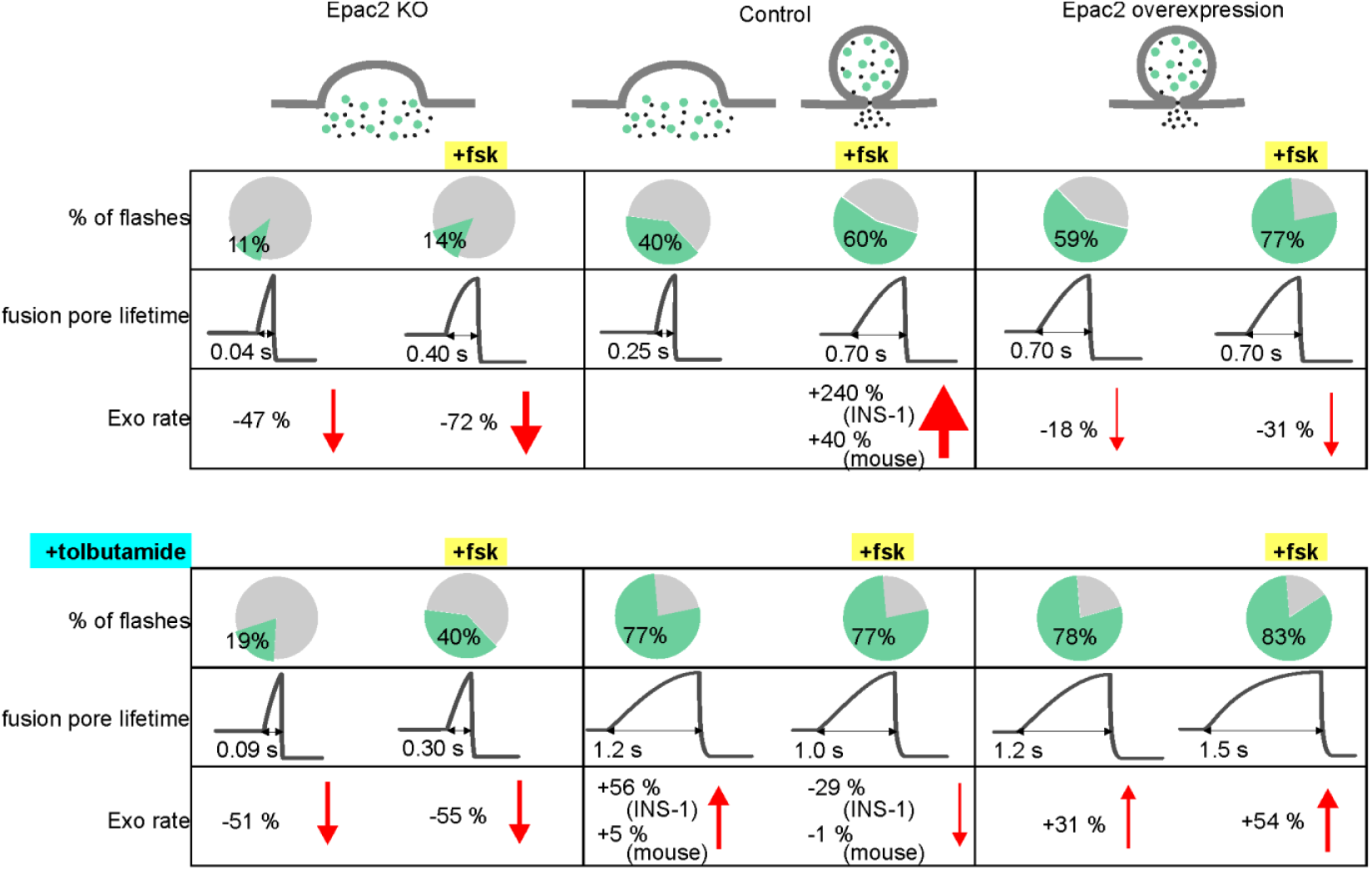
Summary of fusion pore characteristics. Fraction of events with restricted fusion pores, fusion pore lifetime and exocytosis rate for Epac2 KO (first column), controls (second column) and with Epac2 overexpression (third column) in absence (upper rows) and presence of tolbutamide (bottom rows). Changes in exocytosis are compared to controls without (left half columns) or with (right half columns) forskolin.

Paradoxically, two clinically important classes of antidiabetic drugs, GLP-1 analogs and sulfonylureas, activate Epac in β-cells and caused restriction of the fusion pore. Sulfonylureas have long been known to stimulate insulin secretion by binding to SUR1, which results in closure of K_ATP_ channels and depolarization ^37^. The drugs also accelerate PKA-independent granule priming in β-cells, which may involve activation of intracellularly localized SUR1 ^53^. Our data indicate that sulfonylureas exert a third mode of action that leads to the restriction of the fusion pore and therefore limits insulin release. Two pieces of evidence suggest that SUR1 is not involved in the latter. First, acute exposure to sulfonylureas had no effect on fusion pore behavior, although it blocks K_ATP_ channels (indicating SUR1 activation). Only long-term exposure to sulfonylurea resulted in restricted fusion pores, likely because it allowed the drugs to enter the cytoplasm. Second, we could not detect enrichment of SUR1 at the granule release site, which precludes any direct role of the protein in fusion pore regulation. Sulfonylurea compounds have been shown to allosterically stabilize the cAMP-dependent activation of Epac ^54,55^. Our finding that sulfonylurea caused fusion pore restriction in the absence of forskolin indicates that basal cAMP concentrations are sufficient for this effect. It can further be speculated that the competing stimulatory (via exocytosis) and inhibitor effects (via the fusion pore) of sulfonylureas on insulin secretion, contribute to the reduction in sulfonylurea effectiveness with time of treatment. Moreover, long term treatment with GLP-1 analogs disturbs glucose homeostasis ^56^, and combination therapy of sulfonylurea and DPP4 inhibitors (that elevate cAMP) has been shown to lead to severe hypoglycemia ^57^, an effect that likely depends on Epac ^58^.

Epac mediates the PKA-independent stimulation of exocytosis by cAMP ^59^. This effect is rapid ^53^, suggesting that Epac may be preassembled at the site of the secretory machinery. Indeed, Epac concentrates at sites of docked insulin granules ^39^, and forms functionally relevant complexes with the tethering proteins Rim2 and Piccolo ^60,61^. However, the amount of Epac2 present at individual release sites did not correlate with fusion pore behavior, which may indicate that the protein acts indirectly by activating or recruiting other proteins. Indeed, we show here that recruitment of two other proteins, dynamin and amisyn, depends on cAMP. Another known target of Epac is the small GTPase Rap1, for which Epac is a guanine nucleotide exchange factor (GEF). Rap1 is expressed on insulin granules and affects insulin secretion both directly ^62^, and by promoting intracellular Ca^2+^-release following phospholipase-C activation ^63^. By altering local PIP_2_ levels, the latter could affect both exocytosis through C2 domain proteins such as Munc13 ^64^, and PIP_2_-dependent recruitment of dynamin to the plasma membrane ^65^.

An unresolved question is whether pore behavior is controlled by mechanisms that promote pore dilation, or that instead prevent it. Dynamin causes vesicle fission during clathrin-dependent endocytosis ^66^, and since dynamin is present at the exocytosis site and required for the kiss&run mode ^29,30,67^, it may have a similar role during transient exocytosis. An active scission mechanism is also suggested by the finding that granules loose some of their membrane proteins during transient exocytosis ^68^. Capacitance measurements have shown that fusion pores initially flicker with conductances similar to those of large ion channels, before expanding irreversibly ^69^. This could result from pores that are initially stabilized through unknown protein interactions and that eventually give way to uncontrolled expansion. However, scission mechanisms involving dynamin can act even when the pore has dilated considerably beyond limit of reversible flicker behavior ^4,14,51^, and even relatively large granules retain their size during fusion-fission cycles ^6,8^. Separate mechanisms may therefore operate, one that prevents pore dilation by actively causing scission, similar to the role of dynamins in endocytosis, and another by shifting the equilibrium between the open and closed states of the initial fusion pore. Curvature-sensitive proteins are particularly attractive for such roles since they could accumulate at the neck of the fused granule; such ring-like assemblies that have indeed been observed for the Ca^2+^-sensor synaptotagmin ^70^. Active pore dilation has also been proposed to be driven by crowding of SNARE proteins ^71^ and α-synuclein ^36^.

Insulin granules contain millimolar concentrations of a variety of small transmitter molecules (serotonin, ATP, GABA) and ions (zinc, calcium) that, once released, result in biologically meaningful signals within the islet ^72^. GABA release activates both autocrine and paracrine signaling in islets. Activation of GABA-A Cl^-^-channels in α-cells inhibits glucagon secretion ^73,74^, whereas depolarization in β-cells leads to enhanced insulin secretion ^72,75^. Tonic GABA signaling is important for the maintenance of β-cell mass, by promoting α-to-β-like cell conversion ^22,75^ and by suppressing T-cell dependent autoimmune responses ^75,76^. Changes in the local release of GABA could therefore play an important role in the development of type-1 diabetes. Interestingly, not all of a β-cell’s granules contain GABA ^77^, either due to differential expression of GABA transporters or, more likely, kiss&run exocytosis that results in granules without GABA content. The latter effect is well-documented in chromaffin cells, where catecholamine content varies considerably amongst granules ^78^. The release of ATP and other nucleotides from β-cells acts as autocrine positive feedback signal, leading to depolarization and intracellular Ca^2+^-release that result in enhanced insulin secretion ^79–81^, but also negative effects have been reported ^16,17^. Paracrine ATP signaling coordinates Ca^2+^ signaling among β-cells ^15^. In dogs, purinergic signaling stimulates secretion of somatostatin from δ-cells ^82^. Other likely targets of ATP signaling are the islet vasculature, and islet-resident macrophages as part of the immune system ^20^. The latter may play a role for the autoimmune attack in type-1 diabetes, the pro-inflammatory milieu associated with type-2 diabetes ^83^, and β-cell development during embryogenesis ^84^. Finally, human β-cells also secrete serotonin (5-hydroxytryptamine), which inhibits glucagon secretion from α-cells ^85^. The effects of Epac activation by cAMP and sulfonylureas on fusion pore behavior are therefore likely to have profound consequences for intra-islet signaling.

## Acknowledgement

We thank J. Saras, P.-E. Lund, and A. Thonig (Uppsala University) for expert technical assistance, and D. Machado (University of La Laguna) for spike analysis software. The work was supported by the Swedish Research Council, Diabetes Wellness Network Sweden, Swedish Diabetes Society, European Foundation for the Study of Diabetes, Swedish Society for Medical Research, Hjärnfonden, and the NovoNordisk and Family Ernfors foundations. N.R.G. was supported by the European Foundation for the Study of Diabetes (EFSD)/Lilly Research Fellowship and the Swedish Society for Medical Research (SSMF). SD was supported by grants from the Norwegian Research Council (NFR) and Helse-Bergen. Human islets for research were provided by the Nordic Network for Islet Transplantation (supported by JDRF grant 31-2008-416, ECIT Islet for Basic Research Program).

## Author contributions

A.G., N.R.G. and S.B. designed experiments and analyzed the data. A.G. performed imaging and electrophysiology experiments in human β-cells, INS-1 cells, and Epac2^-/-^ β-cells. N.R.G. performed imaging experiments in human and Epac2^-/-^ β-cells. M.O.H. designed and generated the NPY tdOrange virus construct. M.B. and S.D. created the Epac2^-/-^ mice; A.T. maintained the animals and provided islets and critical reagents. S.B. conceived the study. S.B. and A.G. wrote, and all authors critically reviewed the manuscript.

## Online Methods

### Cells

Human islets were obtained from the Nordic Network for Clinical Islet Transplantation Uppsala ^86^ under full ethical clearance (Uppsala Regional Ethics Board 2006/348) and with written informed consent. Isolated islets were cultured free-floating in sterile dishes in CMRL 1066 culture medium containing 5.5 mM glucose, 10% fetal calf serum, 2 mM L-glutamine, streptomycin (100U/ml), and penicillin (100U/ml) at 37°C in an atmosphere of 5% CO_2_ up to two weeks. Prior to imaging, islets were dispersed into single cells by gentle agitation using Ca^2+^-free cell dissociation buffer (Thermo Fisher Scientific) supplemented with 10% (v/v) trypsin (0.05% Thermo Fisher Scientific). INS1-cells clone 832/13 ^87^ were maintained in RPMI 1640 (Invitrogen) with 10 mM glucose, 10% fetal bovine serum, streptomycin (100 U/ml), penicillin (100 U/ml), Sodium pyruvate (1 mM), and 2-mercaptoethanol (50 μM).

Mouse islets were obtained from 5-12 months old WT and Epac2^-/-^ ^45^ with collagenase digestion. Briefly, pancreas of animals was dissected out and fat and connective tissue were removed on ice in Ca5 solution (in mM 125 NaCl, 5KCl, 1.2 MgCl_2_, 1.28 CaCl_2_, 10 HEPES; pH 7.4 with NaOH). Pancreas was injected with Collagenase P (1 mg/ml) and cut into small pieces before mechanical dissociation (7 min at 37 °C). BSA was added immediately and islets were washed 3X with ice cold Ca5 with BSA. Islets were dispersed into single cells using Ca^2+^-free cell dissociation buffer (supplemented with 10% (v/v) trypsin) and gentle agitation. Dispersed cells were sedimented by centrifugation, resuspended in RPMI 1640 medium (containing 5.5 mM glucose, 10% fetal calf serum, 100 U/ml penicillin and 100 U/ml streptomycin).

The cells were plated onto 22-mm polylysine-coated coverslips, and were transduced the next day using adenovirus (human & mouse cells) or transfected the same day with plasmids (INS1 cells, using Lipofectamine2000, Invitrogen) encoding the granule markers NPY-Venus, NPY-EGFP or NPY-tdOrange. Imaging proceeded 24-36 hours later.

### Constructs

The open reading frame of human amisyn (NM_001351940.1) was obtained as a synthetic DNA fragment (Eurofins, Germany) and was cloned into pCherry2 C1 (Addgene, plasmid nr 54563) by seamless PCR cloning. The linker between Cherry2 and amisyn translates into the peptide SGLRSRAQASNSAV. The plasmid N1 NPY-EGFP-mCherry coding for NPY-linker(TVPRARDPPVAT)-EGFP-linker(KRSGGSGGSGGS)-mCherry was made by seamless PCR cloning. The correct open reading frame of both Cherry2-linker-amisyn and NPY-EGFP-mCherry was confirmed by Sanger sequencing (Eurofins, Germany). The NPY-tdOrange2 adeno virus was made using the RAPAd vector system (Cell Biolabs, San Diego, USA). NPY-tdOrange2 ^41^ was cloned into the pacAd5 CMVK-NpA Shuttle plasmid (Cell Biolabs). Virus was produced in HEK293 cells and isolated according to the instructions of the manufacturer (Cell Biolabs).

### Solutions

Cells were imaged in (mM) 138 NaCl, 5.6 KCl, 1.2 MgCl_2_, 2.6 CaCl_2_, 10 D-glucose 5 HEPES (pH 7.4 with NaOH) at 32-34 °C. Exocytosis was evoked with high 75 mM K^+^ (equimolarly replacing Na^+^), applied by computer-timed local pressure ejection through a pulled glass capillary. For K^+^-induced exocytosis, spontaneous depolarizations were prevented with 200 µM diazoxide (50 µM for Fig 5e-h). In Figs 1,2,4-6, forskolin (Fsk; 2 µM) was included. In Fig 1, exendin-4 (10 nM) was included. In Fig 5e-h, exocytosis was evoked by sulfonylureas (500 µM tolbutamide, 200 µM glibenclamide or 200 µM gliclizide). For electrophysiology, glucose was reduced to 3 mM, and the electrodes were filled with (mM) 125 CsCl, 10 NaCl, 1.2 MgCl_2_, 5 EGTA, 4 CaCl_2_, 3 Mg-ATP, 0.1 cAMP, 10 HEPES (pH 7.15 using CsOH).

### TIRF microscopy

Human cells were imaged using a lens-type total internal reflection (TIRF) microscope, based on an AxioObserver Z1 with a 100x/1.45 objective (Carl Zeiss). TIRF illumination with a calculated decay constant of ∼100 nm was created using two DPSS lasers at 491 and 561 nm (Cobolt, Stockholm, Sweden) that passed through a cleanup filter (zet405/488/561/640x, Chroma) and was controlled with an acousto-optical tunable filter (AA-Opto, France). Excitation and emission light were separated using a beamsplitter (ZT405/488/561/640rpc, Chroma) and the emission light chromatically separated (QuadView, Roper) onto separate areas of an EMCCD camera (QuantEM 512SC, Roper) with a cutoff at 565 nm (565dcxr, Chroma) and emission filters (ET525/50m and 600/50m, Chroma). Scaling was 160 nm per pixel.

INS1 and mouse cells were imaged using a custom-built lens-type TIRF microscope based on an AxioObserver D1 microscope and a 100x/1.45 NA objective (Carl Zeiss). Excitation was from two DPSS lasers at 473 nm and 561 nm (Cobolt), controlled with an acousto-optical tunable filter (AOTF, AA-Opto) and using dichroic Di01-R488/561 (Semrock). The emission light was separated onto the two halves of a 16-bit EMCCD camera (Roper Cascade 512B, gain setting at 3,800 a.u. throughout) using an image splitter (DualView, Photometrics) with ET525/50m and 600/50m emission filters (Chroma). Scaling was 100 nm per pixel for INS-1 experiments and 160 nm for mouse cells.

### Image analysis

Exocytosis events were identified manually based on the characteristic rapid loss of the granule marker fluorescence (1-2 frames). Events were classified as flash events if they exhibited apparent increase in the fluorescence signal before the rapid loss of the granule fluorescence. The fusion pore lifetimes were obtained for both types of events by non-linear fitting with a discontinuous function in Origin as described previously ^41^. Protein binding to the release site (ΔF/S) was measured as described previously ^88^.

### Electrophysiology

ATP release was measured in INS1 cells expressing RFP-tagged P2X_2_ receptor ^4^. Cells were voltage-clamped in whole-cell mode using an EPC-9 amplifier and PatchMaster software (Heka Elektronik, Lambrecht, Germany) with patch-clamp electrodes pulled from borosilicate glass capillaries that were coated with Sylgard close to the tips, and fire-polished (resistance 2-4 MΩ). The free [Ca^2+^] was calculated to be 600 nM (WEBMAXC standard) and elicited exocytosis that was detected as P2X_2_-dependent inward current spikes. Currents were filtered at 1 kHz and sampled at 5 kHz. Spike analysis was performed using automated program for amperometric recordings in IGOR Pro ^89^, with the threshold set at eight times the RMS noise during event-free section of recording.

### Statistics

Data are presented as mean ± SEM unless otherwise stated. Statistical significance was tested (unless otherwise stated) using Students t-test and is indicated by asterisks (*p < 0.05, **p < 0.01, ***p < 0.001). Ratios of flash events were tested using Mann-Whitney u-test and fusion pore lifetimes were tested with Kolmogorov-Smirnov test.

